# Individual differences reveal similarities in serial dependencies across perceptual tasks, but no relation to serial dependencies for oculomotor behavior

**DOI:** 10.1101/2024.02.14.580238

**Authors:** Shuchen Guan, Alexander Goettker

**Affiliations:** Justus Liebig Universität Giessen, Germany; Center for Mind, Brain and Behavior, Universities of Marburg, Giessen, and Darmstadt

## Abstract

Serial dependence effects from one trial to the next have been observed across a wide range of perceptual tasks, as well as for oculomotor behavior. This opens up the question of whether the effects observed across all of these studies share underlying mechanisms. Here we measured the same group of observers across four different tasks, two perceptual (color judgments and orientation judgments) and two oculomotor (tracking of moving targets and the pupil light reflex). On the group level, we observed significant attractive serial dependence effects for all tasks, except the pupil response. The rare absence of a serial dependence effect for the reflex like pupil light response suggests that sequential effects require cortical processing or even higher-level cognition. In the following step, we leveraged reliable individual differences between observers in the other tasks to test whether there is a trait-like behavior of some observers showing stronger serial dependence effects across all of these tasks. We observed a significant relationship in the strength of serial dependence for the two perceptual experiments, but no relation between the perceptual tasks and oculomotor behavior. This indicates, differences in processing between perception and oculomotor control and the absence of a general trait-like behavior that affects all tasks similarly. However, the shared variance in the strength of serial dependence effects across different perceptual tasks indicates the importance of a similar positive decision bias present, that is reliably different between observers and consistent across different serial dependence tasks.

## Main Text

A common experimental assumption is that behavioral responses across trials are independent of each other and are only driven by the current sensory input. However, there is converging evidence demonstrating that previous trials systematically influence current responses. Previous experiences often have an attractive effect on the current perceptual judgment, a phenomenon often referred to as serial dependence (see (1, 2) for recent overviews). For example, seeing a leftward-oriented grating in the previous trial will lead to a more leftward response in the current trial (3). Similar attractive sequential effects have been reported across the whole perceptual hierarchy, ranging from simple orientation or color judgments to judgments to complex motion patterns or facial identity (4), but also in attention and memory research (5) or oculomotor control (6). One of the main theories of why these effects occur is that since our natural sensory input contains strong autocorrelations over time, integrating previous information can be beneficial to reduce noise and stabilize perception (3). Interestingly, while this integration seems beneficial, the weight given to past information reliably differs across individuals (7, 8). Together with serial dependence occurring across so many different tasks, this opens up an interesting question: Is the weight given to past information comparable across tasks? Or in other words, is there one person who consistently shows strong serial dependence effects in each task, potentially even similar to a personality trait? By leveraging individual differences across different serial dependence tasks, we show that the strength of sequential effects is correlated across different perceptual tasks, but not with the strength of sequential effects for oculomotor control. These results shed new light on the potential origin of perceptual sequential effects and differences in sensory processing between perception and oculomotor control.

To search for common underlying mechanisms for sequential effects, we measured serial dependencies in four different tasks for the same group of observers (Figure 1A). We used modified versions of three already established tasks: color judgments, orientation judgments, and oculomotor tracking. Additionally, we included a novel domain for oculomotor behavior: the pupil light reflex (9). All tasks had a similar structure; each trial consisted of a prior and a probe stimulus. During the prior, observers saw one of two stimuli that differed across the relevant dimension of the task (e.g. two different orientations for the orientation task or to different speeds for the tracking task). This was then immediately followed by the probe stimulus which was always identical (see Methods for details). This allowed us to make a quick estimate of the strength of serial dependence effects, by looking at the difference in behavior for the probe stimulus depending on which prior stimulus was presented.

**Figure 1.**
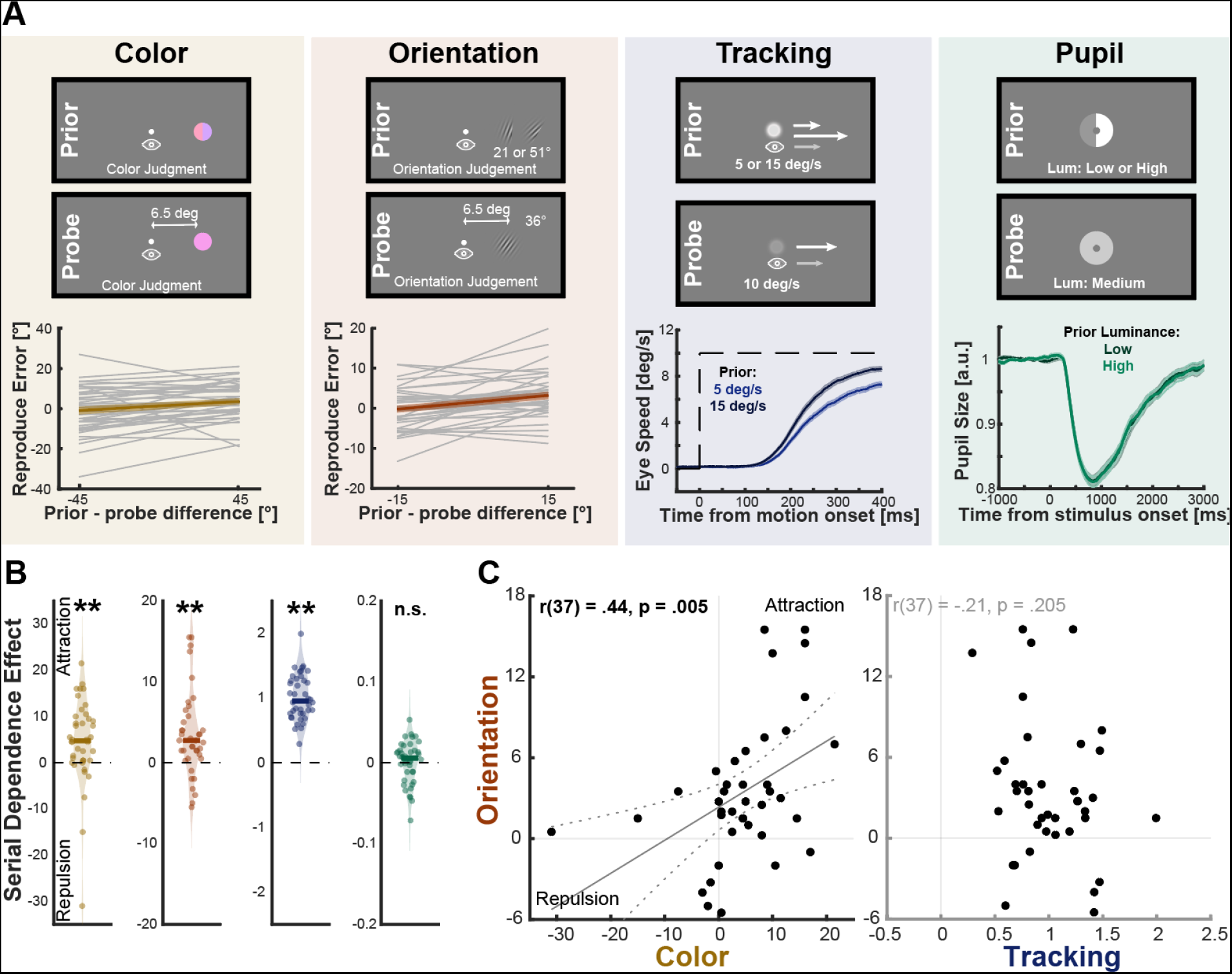
Serial dependence effects across tasks. **A** Depiction of the four different tasks for color judgments, orientation judgments, oculomotor tracking, and the pupil light response. The top rows show a depiction of the paradigms: In the prior always one of two different stimuli was presented. Stimuli were adjusted for visualization, please see Methods for more details. Please note, that while always both prior possibilities are depicted, observers in the tasks always just saw one. The probe stimulus was always identical and in the middle of the two prior stimuli for the relevant dimension. This design allowed us to look at the responses in the probe stimulus with respect to the different priors. The average responses for the probe stimulus separated by the two different priors are shown in the bottom row. **B** The strength of the serial dependence effects is quantified as the differences observed in the probe responses. Individual dots depict individual observers, the solid line depicts the median and the shaded area in the background depicts a violin plot based on a distribution estimate. ** Depict a significant effect on the .01 level. **C** Correlations between the individual serial dependence effects across two tasks. The left panel shows the correlation between the color and orientation task, and the right panel shows the correlation between orientation and oculomotor tracking. Each dot represents one observer, the gray line a linear regression model fitted to the data, and the gray dashed line the 95% CI of the fit.

For the already established tasks, we replicated previous results across the population with our short tasks (Figure 1B). We found significant attractive serial dependence effects for color judgments (BF10 = 11.160), orientation judgments (BF10 = 138.961), and tracking behavior (BF10 = 3.125*10^17^). Despite the significant effects for the whole group of observers, there was substantial variability between observers. For perceptual judgments, some observers showed repulsive effects and for all tasks, the magnitude of serial dependence effects could differ by multiple orders of magnitude across observers. These variations were not just measurement noise but were reliable differences between observers (estimated via split-half Pearson correlation: Color: r = .627; Orientation: r = .493; Oculomotor: r = .344, see Figure S1A & Methods for more Details).

The individual variability in the strength of serial dependence effects was not related to individual differences in basic sensory processes in terms of processing time or response accuracy. The strength of serial dependence effects for each observer was not related to differences between the presentation of the stimuli due to different response durations (Color: r(38) = .193, p = .233; Orientation: r(37) = -.146, p = .376; Pearson correlation, the timing was between stimuli was fixed in the oculomotor task; Figure S1B). In addition, previous work has shown that across trials serial dependence followed a Bayesian weighting with stronger effects for more reliable priors (10). Therefore, we tested whether there is a relationship between the accuracy of the responses for the prior stimulus as a proxy for the reliability and the strength of serial dependence effects. However, neither for the perceptual tasks (Color: r(38) = -.281, p = .079; Orientation: r(37) = .189, p = .250, Pearson correlation) nor for the oculomotor tracking task (r(38) = .153, p = .347; Figure S1C), a systematic relationship was present. This highlights that the variance across observers seems to be related to individual differences in the weighting of previous and current information that cannot be simply explained by differences in the quality of sensory information.

In contrast to the overwhelming literature on sequential effects across all kinds of tasks, for the pupil response, we did not observe a serial dependence effect. There was even significant evidence for the absence of an effect (BF01 = 5.861). The lack of a systematic difference was also supported by the absence of reliable individual differences (r = -.065, see Figure S1A). One could argue, that this null effect can be explained by the relatively long trial duration between the prior and probe stimulus (7s) for the pupil task to give the pupil time to go back to baseline size before the probe occurred, however, we know from a recent meta-analysis that serial dependence effects can last up to 15 s (4). In addition, due to observers also giving a perceptual response for the prior stimulus, the time between the prior and probe stimulus for the perceptual tasks was also quite long (Average Time: 10.63 ± 1.98s for Color; 7.38 ± 2.20s for Orientation). Therefore, we hypothesize that there are two reasons for the absence of sequential effects for the pupil response. First, while small biases in perceptual responses are not harmful, an overdilation of the pupil due to a sequential effect could even potentially damage photoreceptors. Second, the pupil light response is controlled in a reflex-like manner mostly by the brain stem (9). Together with previous work showing that sequential effects for oculomotor control seem to be controlled by frontal brain areas (10), this suggests the need for cortical processing to find sequential effects.

For our main aim of searching for similarities in the strength of serial dependence across tasks, we harnessed the reliable individual differences observed for color and orientation judgments and oculomotor tracking (Figure 1C). We observed that the magnitude of the serial effects was correlated between the two perceptual tasks (r(37) = .439, p = .005). This result complements recent work showing that individual differences within a single task can be predicted by a different weighting of a positive choice and a repulsive motor bias (8). However, they extend our current knowledge by showing that this mapping generalizes across different perceptual tasks. This indicates that there is a substantial amount of shared variability across our two perceptual tasks (20% of the variance), which could point to the role of an individual trait of how strongly previous experience affects the current judgment. In contrast to the two perceptual tasks, there was no systematic relationship between the sequential effects for the two perceptual tasks and oculomotor tracking (r(38) = -.234., p = .147 with color; r(37) = -.208, p = .205 with orientation).

These results highlight two important aspects: First, the lack of a link between perceptual judgments and oculomotor control highlights that information processing for perception and oculomotor control can differ (11). For example, while the mediating signal for perceptual serial dependence effects originates at higher-level visual processing (12), serial dependence effects for oculomotor control seem to rely on low-level retinal information (6). Second, similarities in the strength of the sequential effects across different perceptual tasks provide an exciting starting point: They point to the importance of a more general factor determining the weight of past information across different perceptual tasks. Despite the differences in processing for oculomotor control and perception, the lack of a correlation between the perceptual tasks and the tracking task seems to make it unlikely that the general factor is just related to the sensory integration of past information. It rather points to a more high-level factor that could reflect a common positive decision bias (8), that introduces similarities in serial dependence effects across different perceptual tasks. The next critical step will be to investigate how far this generalization holds across different and more complex tasks: our two perceptual tasks were both related to low-level sensory signals that share some processing resources (13, 14) and required a similar judgment response. Therefore, our two tasks might be inherently more similar to each other than to a task requiring judging facial emotion. However, if this generalization indeed holds across the visual processing hierarchy, it will allow important insights into the shared origins and mechanisms that can explain sequential effects across tasks.

Together, our results highlight the potential of individual differences to gain insights into sensorimotor processing. Leveraging the variations between observers revealed a common underlying explanation that could unify parts of the vast literature on serial dependence effects across the whole visual processing hierarchy. In addition, we found differences between how sensory information is processed for perception and oculomotor control and a rare exception: one of the most basic reflexes of sensorimotor control, the pupil light reflex, did not show serial dependence effects. This indicates that sequential effects require cortical processing.

## Acknowledgments

The authors want to thank Lorena Klöckner for her help with the data collection. S.G. was supported by European Research Council (ERC Advanced Grant Color 3.0, 884116). A.G. was supported by the Supported by the Deutsche Forschungsgemeinschaft (DFG; project number 222641018–SFB/TRR 135 Project A1).

## Author contributions

Conceptualization, S.G. & A.G.; Methodology, S.G. & A.G.; Formal Analysis, S.G. & A.G.; Writing – Original Draft, A.G.; Writing – Review & Editing, S.G. & A.G.; Visualization, S.G. & A.G.; Funding Acquisition, S.G. & A.G.

## Supplemental Results

**Figure S1.**
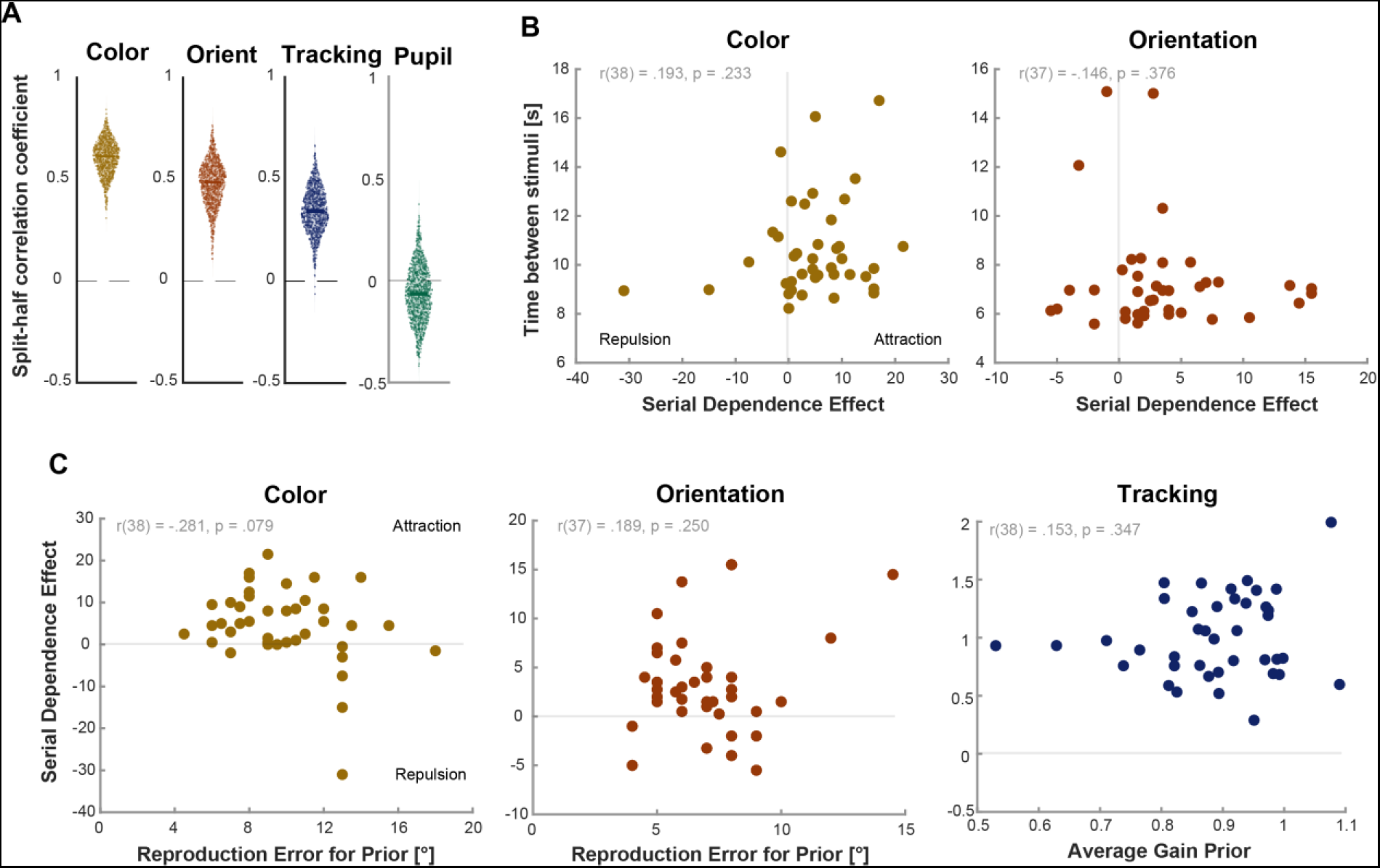
Individual variability in serial dependence effect. **A** Reliability of individual differences through split-half correlation. Each plot displays the distribution of 1000 split-half correlation coefficients for the respective task, represented by 1000 dots. For each correlation, trials were randomly divided into two equal groups for each observer, ensuring equal representation of each prior across groups. The serial dependence effect was calculated separately for each group per observer, followed by the computation of Pearson correlations between the groups’ serial dependence effects across observers. The median coefficient is depicted by a solid line, while the shaded area represents a violin plot, illustrating the distribution estimate. **B** Correlation between individual serial dependence effects and stimulus timing. The left panel illustrates the Pearson correlation between the individual color serial dependence effect and the interval between prior and probe stimuli presentations, with the median interval time used for each observer. The right panel illustrates the correlation between the individual orientation serial dependence effect and the interval between stimuli. Each dot represents an observer. On the x-axis, positive values indicate an attractive serial dependence, whereas negative values suggest a repulsive serial dependence. **C** Correlation between individual serial dependence effects and response accuracy. The left panel displays the Pearson correlation between the individual color serial dependence effect and the median error in reproducing prior stimuli hues. The middle panel examines the correlation between the individual orientation effect and the median error in reproducing prior Gabor orientations. The right panel depicts the correlation between the individual serial dependence effect in tracking task and the accuracy of oculomotor behavior. The accuracy for oculomotor behavior was computed as the average pursuit gain for the prior stimuli. Positive values on the y-axis reflect an attractive serial dependence, while negative values denote a repulsive serial dependence.

### Methods

### Observers

Data were collected from 40 observers (29 females, 25.4 ± 4.7 years old). Observers were students or employees of Giessen University, Germany. All observers were naïve to the study. They had normal or corrected-to-normal vision and passed the Ishihara color blindness test ^S1^. Informed consent was obtained from each observer before data collection. All the tasks complied with the Helsinki Declaration and were approved by the local ethics committee (Giessen LEK 2020-0015).

### Setup

A 32-inch LCD monitor (Display++; Cambridge Research Systems, Ltd., Rochester, UK) with a resolution of 1920 × 1080 and a refresh rate of 120 Hz, was used. The chromaticity and luminance values (xyY) for the monitor primaries are as follows: R = [0.642, 0.34, 27.03], G = [0.30, 0.60, 111.6], and B = [0.15, 0.07, 15.67], and white = [0.29, 0.302, 155.2]. The monitor is characterized by linear gammas (R: 1.02, G: 1.04, B: 1.03). Observers were sitting comfortably in a dark room, with their chin and forehead positioned on a rest to ensure stability. Their eyes were aligned with the screen center at a viewing distance of 85 cm. Eye movements from the right eye were tracked using a desk-mounted EyeLink 1000 Plus eye tracker (SR Research, Kanata, ON, Canada) at a sampling rate of 1000 Hz. The tasks were programmed in MATLAB (MathWorks, Natick MA) using the Psychtoolbox^S2^. A nine-point calibration procedure was conducted before each block to precisely align gaze data with the screen coordinates.

### Experimental Paradigm

Four different tasks were measured for each observer (Figure 1): color judgment, orientation judgment, oculomotor tracking, and testing of the pupil light reflex. The sequence of these tasks was randomized across observers. Each observer completed two sessions, with one block dedicated to each task in every session. The sessions were conducted on separate days, each lasting between 1 to 1.5 hours. All tasks shared a similar structure; in each trial, two stimuli were presented: the prior, which was presented first and differed from the probe stimulus in two opposite directions. This was followed by the probe stimulus, which remained constant. By examining the effect of the prior stimulus on the observers’ responses to the probe stimulus, we quantified the serial dependence effect.

### Color judgment

Each trial consisted of a prior and a probe stimulus, following identical procedures. The stimuli were all defined within the DKL color space^S3^, and presented on a gray (at the isoluminant plane of the DKL space, with xyY = [0.29, 0.31, 80.56]) background. The hues of both the prior and probe stimuli were set at the isoluminant plane. For the prior stimulus, two hues were used: an azimuth of 275° (+5° difference from -[S-(L+M)] axis, close to purple, xyY = [0.26, 0.22, 80.36]) and 355° (−5° difference from +[L-M] axis, close to pink, xyY = [0.33, 0.28, 80.60]), while the probe stimulus’s hue was set to the midpoint between the two prior hues, with an azimuth of 315° (close to magenta, xyY = [0.29, 0.23,79.97]).

The trial started with a white fixation dot (0.5 deg in diameter) displayed at the center of the screen for 2s. Subsequently, a colored circle (4 deg in diameter) appeared in the right visual field at 6.5 deg eccentricity for 300 ms. Observers were instructed to maintain their gaze on the dot while using their peripheral vision to perceive the color of the circle. After the circle disappeared, a color mask of the same size emerged at the same location for 1s to erase any potential after-effect. This color mask was a pixelized pattern, with each pixel randomly selected from a range of 0° to 360° hue on the isoluminant plane. The mask then disappeared, and after a short interval (250 ms), a color square (size: 1 × 1 deg) appeared at the center of the screen, surrounded by a full-range color ring (at 7.5 deg eccentricity, width: 1 deg), covering 0-360° hue at the isoluminant plane. Observers were required to adjust the color of the square to match the previously seen color of the target circle. The initial color of the square was randomly selected out of all possibilities. They could make either a big-step change (using Left/Right arrow keys, each step amounting to a 3° change on the isoluminant plane) or a small-step change (using Up/Down arrow keys, each step was 1°), without any time limitation. Following the response period, a blank screen was shown for 200 ms. In each block, every prior stimulus was repeated 20 times, leading observers to complete a total of 40 trials (2 priors x 20 repeats), resulting in 80 judgments (each prior followed by a probe) and observers completed two blocks for a total of 160 judgments.

### Orientation judgment

The orientation judgment procedure paralleled that of the color judgment. Initially, a fixation circle was displayed for 2s, followed by an oriented Gabor patch (spatial frequency: 1/3 cycles/deg, peak contrast: 20%, Gaussian envelope: 1.5 deg s.d.) positioned in the right visual field at 6.5 deg eccentricity for 300 ms. Observers were required to use their peripheral vision to determine the orientation of the Gabor. Subsequently, a 1/f noise mask embedded in the same Gaussian envelope was presented for 1s. During the response period, an adjustable white line (length: 3.6 deg, width: 3 pixels) appeared at the same location. The initial orientation of the square was randomly selected out of all possibilities. Similar to the color task, observers used the arrow keys to adjust the line’s orientation to match their perceived orientation of the Gabor patch. Upon completing the adjustment, they pressed the space bar to proceed, which either started the same sequence for the probe stimulus or, if the response was made for the probe stimulus, started the next trial.

All stimuli were displayed on a mid-gray background (82.68 cd/m^2^). The probe Gabor was set to a fixed orientation of 36°. The prior was presented with a 15° shift either clockwise (21°) or counter-clockwise (51°). In each block, every prior orientation was repeated 20 times, culminating in a total of 40 trials and yielding 80 judgments. Observers completed two blocks.

### Oculomotor tracking

For the oculomotor task, a single trial consisted of two movements that needed to be tracked by the observer: The prior movement, which could move with either 5 or 15 deg/s and the probe movement which always moved at 10 deg/s. The direction of the movement was randomly either to the left or right, but the prior and probe movement always moved in the same direction. Before the start of a trial, participants saw a red fixation cross and started the trial by looking at it and pressing the space bar. This was step was used as a drift correction. Then a red fixation circle (diameter 0.2 deg) was presented for a random time between 1 and 1.5s. Then the dot disappeared and the target, a Gaussian blob (SD = 0.4 deg) appeared and immediately moved across the screen. The contrast for the prior stimulus was set to 1, whereas the contrast for the probe stimulus was set to 0.1 to maximize a potential serial dependence effect. The target always appeared with a slight offset into the opposite direction of the target movement; the offset was scaled in a way that the target always crossed the initial fixation location after 200 ms (e.g. for a speed of 10 deg/s, the step was 2 deg). This step-ramp paradigm^S4^ was used to reduce the need for an initial corrective saccade. The target then kept moving for 1 s and disappeared. Then a new fixation circle appeared to indicate the start of the probe movement. After a new random time between 1 and 1.5s, the probe movement was presented with the same temporal characteristics as the prior. When the probe movement disappeared, the appearance of a new fixation cross indicated the start of the next trial.

Each of the prior speeds (5 and 15 deg/s) were presented 15 times per block, leading to the tracking of a total of 60 movements (30 priors and 30 probe movements) per block. Again, observers completed a total of two blocks.

### Pupil light reflex task

At the onset, observers underwent a 1-minute adaptation period for a mid-gray blank screen (50% intensity, 81.68 cd/m^2^) within a dark room. Following this initial adaptation phase, pupillary data recording began. Before each trial, observers were presented with a red fixation cross, and they initiated the trial by fixating on the cross and pressing the spacebar. This step was again used as a drift correction. Once the trial began, a black fixation dot (0.3 deg in diameter) appeared for 1s. Subsequently, a darker/brighter grayish prior disk (10 deg in diameter) appeared alongside the fixation dot for 1s, followed by a blank period with fixation lasting 7s for pupillary measurement. Following this, a probe disk (10 deg in diameter) appeared for 1s and transitioned back to a blank screen with fixation for 7s. There were two prior conditions: one with a high luminance disk (100% intensity, 156.2 cd/m^2^), brighter than the probe as a medium luminance disk (80% intensity, 128.4 cd/m^2^), and a low luminance disk (60% intensity, 96.90 cd/m^2^), darker than the probe.

In the pupillary task, observers were instructed to minimize blinking during the trial, with the flexibility to blink before initiating each trial. All frames within the trials, except for the initial 1-minute adaptation, featured a black fixation dot at the center of the screen to mitigate potential location bias in pupil size measurement. Each prior condition was repeated 10 times, resulting in a total of 20 trials for one block. Again, observers completed two blocks.

### Data Analysis & Preprocessing

Please note that all Data (osf.io/psxqh) as well as the analysis and experimental scripts (github.com/AlexanderGoettker/SerialDependenceTasks) will be made publicly available when the manuscript is published in a peer reviewed journal.

### Color Task

The magnitude of the serial dependence effect was quantified by analyzing the hue reproduction errors for the probe (azimuth of 315°) in each trial, calculated as the difference between the reproduced hue azimuth and the probe’s azimuth. These errors were compared across two conditions, corresponding to the presentation of two different priors (with azimuths of 275° and 355°).

For each observer, the median probe reproduction error was calculated for all trials within each prior condition. The difference between these median errors for the two conditions was then calculated to establish a metric for the serial dependence effect. We identified whether an observer exhibited an attractive serial dependence (where probe reproduction errors were more negative with 275° priors than with 355° priors), a repulsive serial dependence (where probe reproduction errors were more positive with 275° priors than with 355° priors), or no effect.

### Orientation Task

The analysis of the orientation task paralleled that of the color task. We calculated the orientation estimation errors for the probe Gabor (36°) for each trial, defined as the difference between the reproduced orientation and the probe’s orientation. These errors were then compared across trials that featured two different prior Gabors (with orientations of 21° and 51°). An attractive serial dependence was identified if the reproduction errors for the probe Gabor were more negative with priors set at -15° than with 15° priors. Conversely, a repulsive serial dependence was suggested if the reproduction errors were less negative or more positive with -15° priors compared to 15° priors. Since we had one observer, who showed very poor accuracy in the orientation judgment task (average error of around 79°) and was more than 3 times the standard deviation away from the mean of the other observers, we excluded him from this part of the analysis.

### Oculomotor Task

Eye movement data saved and analyzed off-line using custom Matlab scripts. First, blinks were linearly interpolated and the eye position was filtered with a second-order Butterworth filter, with a cutoff frequency of 30 Hz. Then eye velocity was calculated as the first derivative of the filtered position traces. Saccades were identified based on the EyeLink criteria with a speed and acceleration threshold of 30 deg/s and 4000 deg/s2, respectively. After the detection of saccades, a linear interpolation of the eye movement velocity around the time of the saccade (from 35 ms before the detected saccade onset to 35 ms after the detected saccade offset) was performed. Eye movement velocity was filtered with an additional low-pass Butterworth filter with a cutoff frequency of 20 Hz.

Since all targets moved horizontally, we took the horizontal velocity of the eye and aligned it to the target movement onset. To ensure that there was a valid tracking response, we detected pursuit onset in the prior and probe trials as the first point where the horizontal eye velocity reached 30% of the target speed and stayed there for at least 100 ms. Then, for each observer, across all relevant trials, the median eye velocity trace for the probe stimulus was computed, and this was done separately depending on the prior velocities. The serial dependence effect was then computed as the average difference between the two velocity traces from 100 to 350 ms after target motion onset. A positive effect indicated a faster eye velocity for the faster prior.

To ensure our data quality, we excluded trials from the analysis that had more than 500 ms of missing data in a single trial, where the computed velocity after the interpolation of the saccades still was larger than 30 deg/s (indicating some overlooked saccades), and trials where we couldn’t identify a valid pursuit response with a latency below 400 ms. This was done both for the prior and probe movement and only trials where both movements fulfilled all criteria were used for the analysis. With these criteria, we included a total of 4347 out of 4800 trials (90.5%) in the analysis.

### Pupil Task

The eye movement data were saved during the task and analyzed offline with custom Matlab scripts. First, the pupil size during blinks was linearly interpolated (−100 ms before blink to +100ms after blink) and then was filtered with a second-order Butterworth filter, with a cutoff frequency of 30 Hz. Then we aligned the pupil response to the stimulus onset of the probe movement. To account for individual differences in pupil size, the pupil size was normalized based on the average pupil size 1000 ms before a stimulus was presented (the fixation period before the prior and the probe stimulus was presented). As for the eye velocity, we then computed the median pupil size response to the probe stimulus, separately for each of the prior stimuli. To compute the serial dependence effect, for each observer we computed the mean difference in pupil size in the 1s where the probe stimulus was presented. A positive value indicated a stronger pupil light reflex for the brighter prior.

Trials were excluded from the analysis when we had missing data for more than 500 ms. Based on these criteria we included a total of 1555 out of 1600 trials (97.2%) in the analysis.

### Statistical Procedures

To test for serial dependence effects on the group level for each task, we took the respective metric and computed one-sampled Bayesian t-tests. We tested whether the distribution was significantly different from zero. The strength of the Bayesian tests is that they also allow the interpretation of evidence for the absence of an effect.

To ensure that we observed meaningful individual differences, we estimated the reliability of the individual differences in each of the tasks via a split-half correlation. For this, we used a bootstrapping procedure, where instead of taking all trials to compute the average response for the probe stimulus with a respective prior trial, we randomly split the trials into two groups.

We did this for both priors and then computed the serial dependence effect in the same way as before. To estimate a distribution for each observer, we repeated this procedure 1000 times. This gave us for each observer 1000 estimates of the serial dependence effect for one half of the trials and another 1000 estimates for the second half of the trials. We repeated this procedure for each of the observers and then for each of our 1000 iterations computed the Pearson correlation between the serial dependence effects estimated for one half and the serial dependence effects for the other half across observers. This distribution of 1000 correlations gave us an estimate of the reliability of the individual differences for the respective task.

To look into the potential origins of the individual differences we conducted two control analyses. First, in the color and orientation judgments tasks, the trial durations were variable and dependent on the time it took observers to give a response to the prior stimulus. To assess whether the time between the presentation of the prior and probe stimulus influenced the serial dependence effect, we conducted Pearson correlations between the individual strength of the serial dependence effect and the median duration between the two. Second, we wanted to relate overall differences in task performance to the strength of the serial dependence effect. We hypothesized that overall more accurate observers might rely more on the prior stimulus. For that, we computed a proxy for behavioral accuracy in the prior: for the orientation and color judgment tasks, the median of absolute response errors relative to prior stimuli was used as the accuracy metric for each observer. To compute the accuracy of tracking performance, we computed the average pursuit gain for the prior movement in the interval of 100 to 300 ms after pursuit onset. Since due to the lack of a ground truth, we could not compute an accuracy measurement for the pupil response, we did not perform this analysis for the pupil task.

In the last step of the analysis, we correlated the strength of the serial dependence effects across tasks. We did so for each of the tasks that showed a significant serial dependence effect on the group level (Color, Orientation & Tracking).

